# Regulation of FtsZ levels in *Escherichia coli* in slow growth conditions

**DOI:** 10.1101/342766

**Authors:** Jaana Männik, Bryant E. Walker, Jaan Männik

**Affiliations:** Department of Biochemistry & Cellular and Molecular Biology, University of Tennessee, Knoxville, TN 37996, USA; Department of Physics and Astronomy, University of Tennessee, Knoxville, TN 37996, USA

**Author notes:** **Corresponding author:** Jaan Männik.

**Keywords:** cell division, cell cycle regulation, FtsZ, ClpXP protease

## Abstract

A key regulator of cell division in most walled bacteria is the FtsZ protein that assembles into protofilaments attached to the membrane at midcell. These dynamic protofilament assemblies, known as the Z-ring, act as a scaffold for more than two dozen proteins involved in synthesis of septal cell envelopes. What triggers the formation of the Z-ring during the cell cycle is poorly understood. In *Escherichia coli* model organism, the common view is that FtsZ concentration is constant throughout its doubling time and therefore regulation of assembly should be controlled by some yet to be identified protein-protein interactions. Here we show using quantitative analysis of newly developed fluorescent reporter that FtsZ concentration varies in a cell-cycle dependent manner in slow growth conditions and that upregulation of FtsZ synthesis correlates with the formation of the Z-ring. About 4-fold upregulation of FtsZ synthesis in the first half of the cell cycle is followed by its rapid degradation by ClpXP protease in the last 10% of the cell cycle. The initiation of rapid degradation coincides with dissociation of FtsZ from the septum. Altogether, our data indicate that the Z-ring formation in slow growth conditions in *E. coli* is controlled by a regulatory sequence where upregulation of an essential cell cycle factor is followed by its degradation.

**Significance:** FtsZ is the key regulator for bacterial cell division. It initiates division by forming a dynamic ring-like structure, the Z-ring, at the mid-cell. Here we show that, contrarily to the current paradigm, FtsZ concentration in *Escherichia coli* model organism varies throughout cell cycle in slow growth conditions. Faster FtsZ synthesis in the first half of the cell cycle is followed by its rapid degradation by ClpXP protease in the end of the cell cycle. Upregulation of FtsZ synthesis correlates with the formation of the Z-ring. Our data demonstrates that in slow growth *E. coli* cell division progresses according to paradigm where upregulation of essential cell cycle factor is followed by its degradation.

## Introduction

Cell division in *E. coli* and other bacteria is an essential process subject to spatial and temporal coordination with the chromosome replication cycle (1, 2). The coordination guarantees with high fidelity that bacteria in steady-state growth divide once per initiation of chromosome replication, and that division occurs between completely replicated chromosomes. Significant progress has been made in identifying and unravelling the molecular mechanism involved in correct positioning of the bacterial cell division machinery, the divisome, relative to the chromosomes. Currently, three partially redundant molecular systems are known in *E. coli* that are involved in the positioning of its divisome (3). At the same time it has remained largely unclear how the division process is controlled temporally.

The earliest defined event in bacterial cytokinesis is assembly of the tubulin-like FtsZ protein into a dynamic ring-like polymer structure, the Z-ring, which localizes at midcell (4, 5). FtsZ is a GTPase that is conserved in nearly all walled bacteria (6). In the presence of GTP FtsZ monomers polymerize into dynamic protofilaments (7, 8). It has recently been shown that these filaments treadmill (9-11). The protofilaments form higher order structures such as bundles and sheets *in vitro* (12, 13) and likely also *in vivo* (14). In *E. coli* protofilaments are tethered to the inner membrane via FtsA (15) and ZipA (16) linkers. These two proteins, together with FtsZ and non-essential Z-ring associated proteins ZapA-D, form an early divisome complex, the Z-ring (5). After formation of the Z-ring, but with a distinct delay (17), about 30 other proteins are recruited to the division site in a specific order forming a mature divisome (18, 19). Once divisome has matured, it then carries out synthesis of septal cell wall and membranes (20) and safeguards DNA partitioning between daughter cells (21, 22). FtsZ acts as a central scaffold in recruiting and maintaining the Z-ring throughout cell division except the latest stages of constriction (5). Various intrinsic and environmentally generated signals couple to the FtsZ scaffold and lead to its depolymerization in unfavorable conditions.

Currently, it is not clear what triggers formation of the Z-ring. According to the common view, FtsZ concentration during the cell cycle remains constant in both *E. coli* (1, 23, 24) and in distantly related *B. subtilis* (25). At the same time, the cellular concentration of FtsZ (6-7 µM) is much higher than the concentration needed for assembly of FtsZ protofilaments *in vitro* (1 µM) (8). This has led to a paradigm that assembly of the Z-ring is controlled at the level of its assembly (23, 26) meaning that not protein levels but their interactions inhibit the Z-ring formation despite high concentration of FtsZ. Changes in these interactions reflecting possibly DNA replication status (27) may trigger the formation of the Z-ring. Spatial regulators, such as SlmA mediated nucleoid occlusion (28) and the Ter linkage (29) in *E. coli* and Noc protein in *B. subtilis* (30) could potentially couple DNA replication cycle and Z-ring (31) although experimental evidence to support this idea is lacking.

Even though a constant FtsZ concentration through the cell-cycle is currently the accepted view, numerous earlier reports show that FtsZ mRNA levels oscillate during cell cycle in *E. coli* (32-35). Cell-cycle dependent variations in the levels of FtsZ have been described also in other bacteria, including *Caulobacter crescentus* (36), *Prochlorococcus* sp. (37), and *Synechococcus elongatus* (38). It is not clear how the cells could possibly achieve constant levels of protein product from oscillating levels of transcript taken that ribosome levels during the cell cycle remain approximately constant (39). Recently, 15% variation in FtsZ concentration during the cell cycle was reported in faster growth rates in *E. coli* (40), but the origin of these variations has remained unclear. Here we show using a recently developed FtsZ fluorescent reporter (41) that FtsZ concentration in *E. coli* varies more significantly during the cell cycle as the growth rate decreases. We determine that in slow growth conditions the variation in FtsZ levels is achieved by increasing the synthesis rate in the beginning and the degradation rate at the end of the cell cycle. We show that on average 20% of existing cellular FtsZ is degraded in the end of the cell cycle in slow growth conditions by ClpXP protease. Cell-cycle dependent increase in the concentration and amount of FtsZ correlates with the formation of the Z-ring in slow growth conditions, but it is not a sole factor determining the timing of the Z-ring formation.

## Results

### FtsZ concentration varies during cell cycle in slow growth conditions

Cell cycle dependent variations in FtsZ concentration in *E. coli* have not yet been reported in live cells. Recently, functional FtsZ fluorescent fusions have been developed, which can be expressed from the native locus and are the sole copy of FtsZ (41). Here we use one of these constructs, where mNeonGreen (mNG) fluorescent protein is sandwiched between residues 55 and 56 of FtsZ (FtsZ-mNG), to determine the changes of FtsZ numbers and concentration during the cell cycle (detailed description of all strains is in SI Table S1). For measurements of cells in steady state conditions we grew them in microfluidic mother machine channels (42, 43). The linear colonies in mother machine channels allow determining spatial distribution of fluorescent intensities from individual cells through the division process without uncertainties introduced by nearby cells. Such uncertainties are present when agarose pads and dishes are used for imaging. As a first step in the analysis we collected intensity profiles from time-lapse measurements to kymographs (Fig. 1A, B). These kymographs here and in later sections are extended beyond the division events to bring more clearly out variations of FtsZ levels during the cytokinesis. We studied cells in slow growth conditions in M9 glycerol medium with doubling time of T_d_=160±30 min (Fig. 1A) and in faster growth with doubling time of T_d_=74±19 min (mean±s.d.) where M9 glucose was supplemented with casamino acids (Fig. 1B). Note that all measurements were performed at 28°C. In addition to the strain carrying FtsZ-mNG, we also measured the FtsZ concentration in a control strain that expressed an FtsZ C-terminal fusion to GFP (FtsZ-GFP) under control of *lac-*promoter from a plasmid (SI Fig. S1) (9, 44). The FtsZ-GFP strain shows comparable doubling times in slow T_d_=160±29 min and fast T_d_=80±24 min growth conditions. In the strain with plasmid expressed FtsZ-GFP, one would not expect to observe cell cycle dependent variations in the concentration of the fluorescent proteins. Consistent with this expectation, the concentration of fluorescent FtsZ from plasmid expressed FtsZ-GFP varied less than 10% during the cell cycle in both slow (Fig. 1C) and fast growth conditions (Fig. 1B) in population average measurements. In contrast, we found a 37% variation of endogenously expressed fluorescent FtsZ-mNG in slow growth conditions. In fast growth conditions, however, the variation of FtsZ concentration was essentially at the level of the control (10%). The measurements thus show that FtsZ levels in *E. coli* undergo cell cycle dependent oscillations with an amplitude that depends on growth conditions.

**Figure 1.**
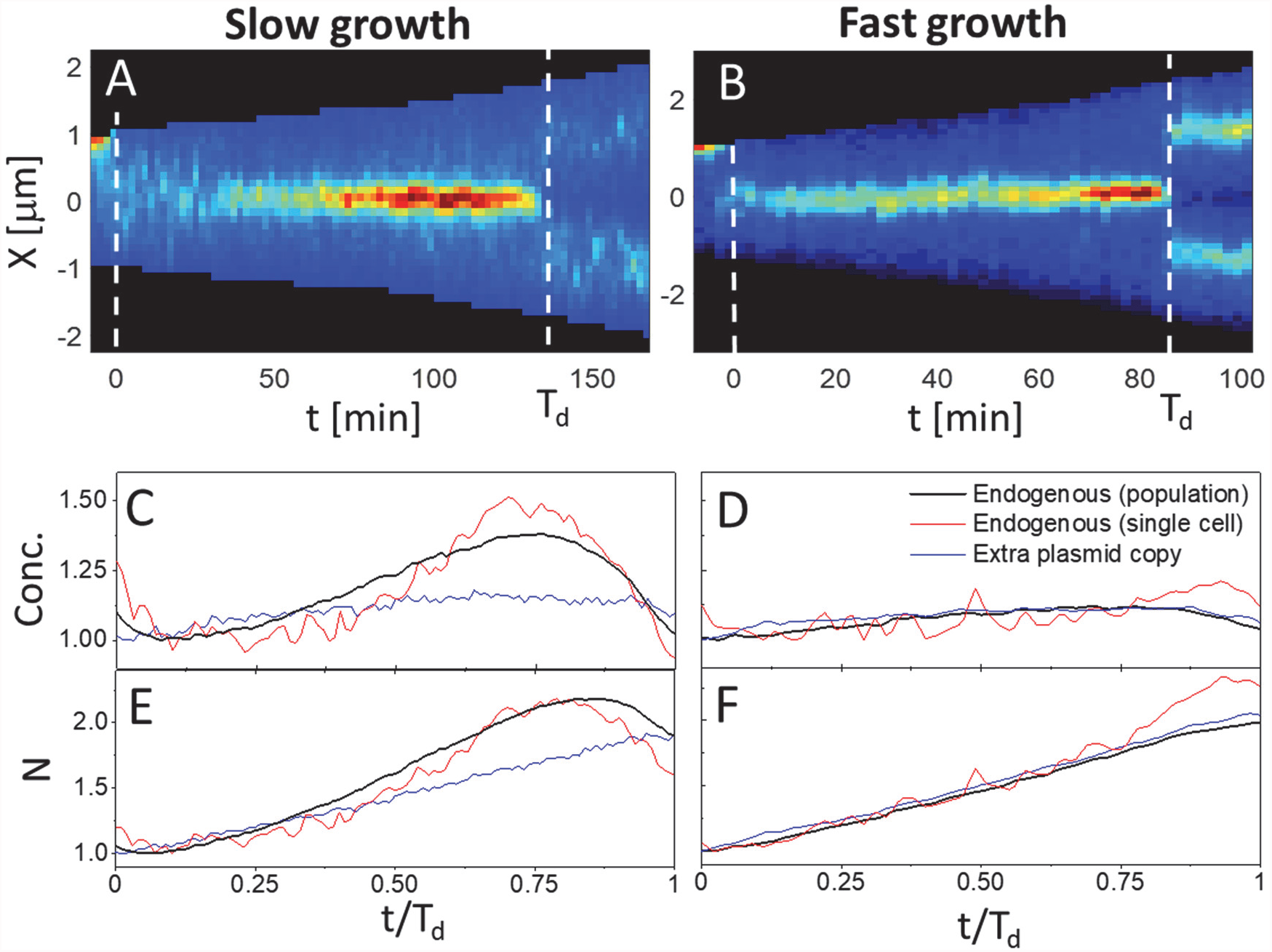
Changes in fluorescent FtsZ concentrations and numbers during cell cycle. Left column corresponds to slow and right column to fast growth conditions. (A, B) Kymographs of FtsZ-mNG intensity distribution along the long axes in representative cells. Red corresponds to high concentration and blue to low. Black marks areas outside the cell. Vertical dashed lines mark times of cell division. (C) Concentration of fluorescent FtsZ construct as a function of cell cycle time in slow growth conditions. Thick black line corresponds to the population average data from the strain expressing a sole FtsZ-mNG from the native locus (N=53), thin blue line to the strain where FtsZ-GFP is expressed from a plasmid in addition to native unlabeled copy (N=16) and thin red line to the cell shown on the kymograph above. (D) The same for fast growth conditions. N=32 for endogenous and N=33 for plasmid expressing strain. (E, F) Numbers of fluorescent FtsZ in the cell in slow and fast growth conditions, respectively. All curves in panels (C)-(F) are normalized by the cell cycle minimum value.

In parallel to the concentration we also determined the variation of fluorescent FtsZ during the cell cycle (Fig. 1E, F). Instead of a monotonic increase, the number of these molecules decreased at the end of the cell cycle in slow growth conditions (Fig. 1E). The decrease started at about 0.9T_d_ when fluorescent FtsZ numbers reached 2.2 times higher level than what is present in the beginning of the cell cycle. The decrease was absent in fast growth conditions (Fig. 1F) and in the control strain with plasmid expressed FtsZ in both growth conditions (Fig. 1E-F, SI Fig. S1). While drop in concentration of FtsZ in the end of the cell cycle could have possibly arisen from slowdown of its synthesis at the end of the cell cycle and concurrent dilution of the protein due to cell growth, the decrease in numbers (about 20%) shows that FtsZ is degraded in the end of the cell cycle. The data furthermore show that the relative amount of FtsZ degraded at the end of cell cycle decreases with the growth rate.

### Degradation in the end of cell cycle is caused by ClpXP

We next investigated what factors could be responsible for the fast degradation of FtsZ in the end of the cell cycle. Earlier studies have identified that FtsZ is one of the substrates for ClpXP protease in *E. coli* (45, 46). The ClpXP protease consists of hexameric ClpX chaperone that unwinds target proteins by passing them via its central pore in ATP-dependent manner and ClpP peptidase that consists of two heptametric rings (47). The identified ClpX binding sites in FtsZ are localized at its C-terminal domain and in its unstructured linker (48). The earlier studies, however, have not revealed changes in ClpXP activity during the cell cycle (46). To test if ClpXP is responsible for the degradation of FtsZ in the end of cell cycle in slow growth conditions, we constructed *ΔclpP* and *ΔclpX* strains in FtsZ-mNG background. Additionally we also constructed *ΔclpA* strain as a control. ClpA in complex with ClpP acts also as a protease (49). However, ClpA, which binds and unwinds target proteins, has so far not been shown to interact with FtsZ in *E. coli*. Concentration of fluorescent FtsZ was essentially constant during the cell cycle in *ΔclpP* and *ΔclpX* strains (Fig. 2A). There was also no apparent degradation in the end of the cell cycle, which could be inferred from the decrease of FtsZ number in these two strains (Fig. 2B). At the same time, the *ΔclpA* control strain behaved very similar to the one without the deletions. From here on, we will refer to the latter as the wild type (WT) strain notwithstanding its modified FtsZ. We found that cell cycle average concentrations and FtsZ levels in *ΔclpP* and *ΔclpX* strains were a factor of 4 higher than in WT (Fig. 2C) while in *ΔclpA* strain the difference was only 1.2 times. The latter was also statistically significant (p<10^-6^; t-test) indicating that ClpAP might have low activity towards FtsZ or have an effect on FtsZ expression. Much higher levels of FtsZ and lack of its cell cycle dependent variation in *ΔclpP* and *ΔclpX* strains provide strong evidence that ClpXP is responsible for degradation of FtsZ in the end of the cell cycle in WT cells.

**Figure 2.**
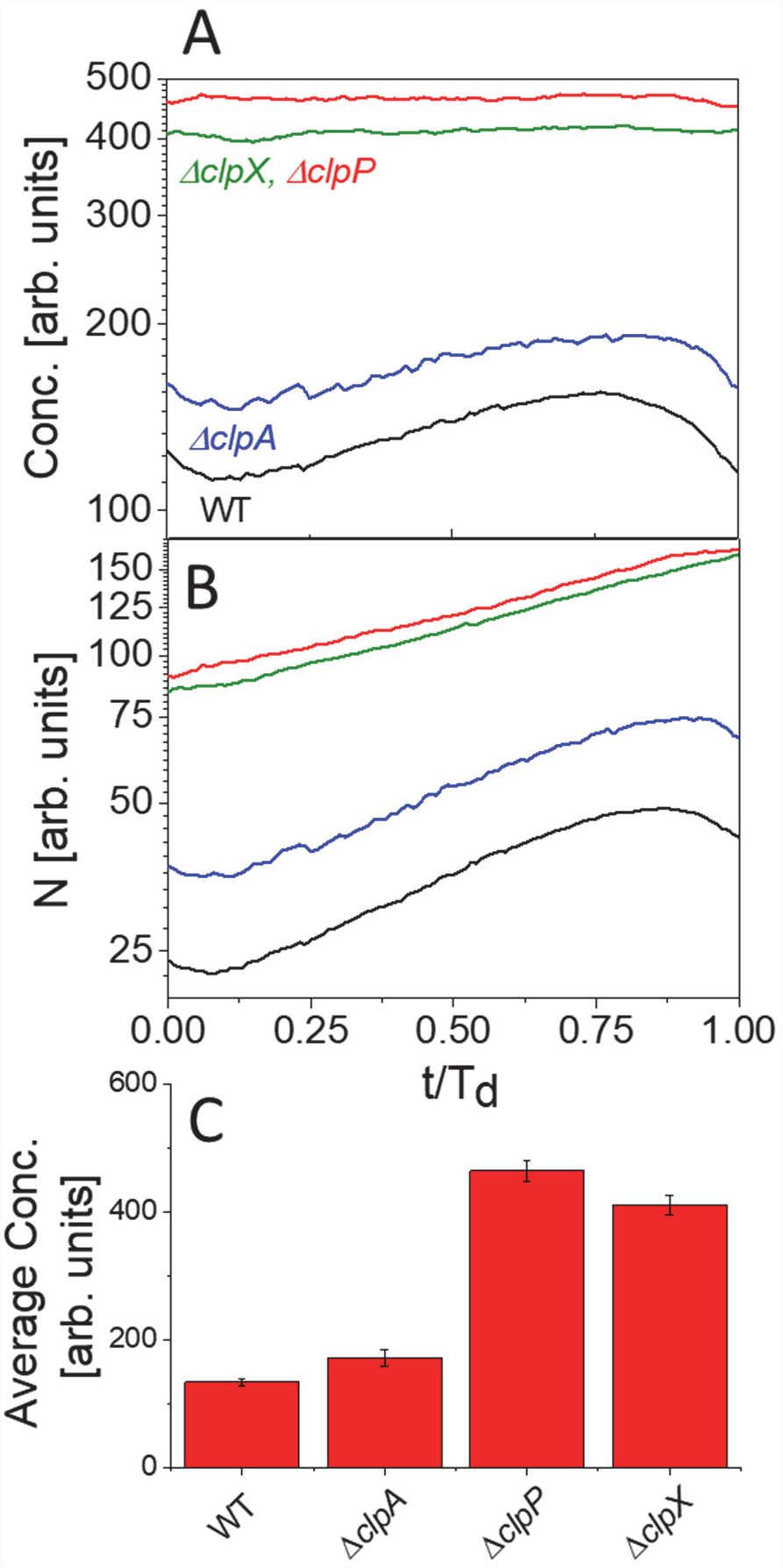
Comparison of FtsZ abundances and concentrations in WT and the protease deletion strains. (A) Population average concentration of FtsZ-mNG as a function of cell cycle time for WT (N=53), Δ*clpA* (N=16), Δ*clpX* (N=31) and Δ*clpP* (N=36) strains. All strains have been measured in slow growth conditions and express FtsZ-mNG from the native locus. (B) Total number of fluorescent FtsZ-mNG in these strains as a function of cell cycle time. Vertical axes in panels (A) and (B) is logarithmic. (C) Concentration of fluorescent FtsZ-mNG averaged over cell cycle in different strains. Error bars are s.e.m.

### Synthesis of FtsZ during the cell cycle is not uniform in slow growth conditions

Closer inspection of data from individual *ΔclpP* and *ΔclpX* cells reveals that FtsZ numbers do not increase uniformly in time (Fig. 3A,B) as they do in population-averaged data (cf. Fig. 2A). The time varying rate of change of FtsZ numbers indicates that synthesis of FtsZ is cell cycle dependent in slow growth conditions. In *ΔclpP* and *ΔclpX* cells we can distinguish two periods in the cell cycle that have different FtsZ synthesis rates. These periods correspond to two different slopes in N vs t curves (Fig. 3B). To find the beginning of these periods and rate of change in FtsZ numbers in each period we performed a piecewise linear fit to the curves from individual cells. Compilation of the fitting results shows (Fig. 3C) a period of upregulated FtsZ synthesis during the cell cycle that starts at time T_1_=0.35±0.2T_d_ and runs until time T_2_=0.85±0.15T_d_ (mean±s.d.). During this period the synthesis rate is about three times higher than that during the rest of the cell cycle. Large variations in start and end times of this period among cell population explain why in population averaged data (Fig. 2A) the period with higher synthesis rate did not reveal itself.

**Figure 3.**
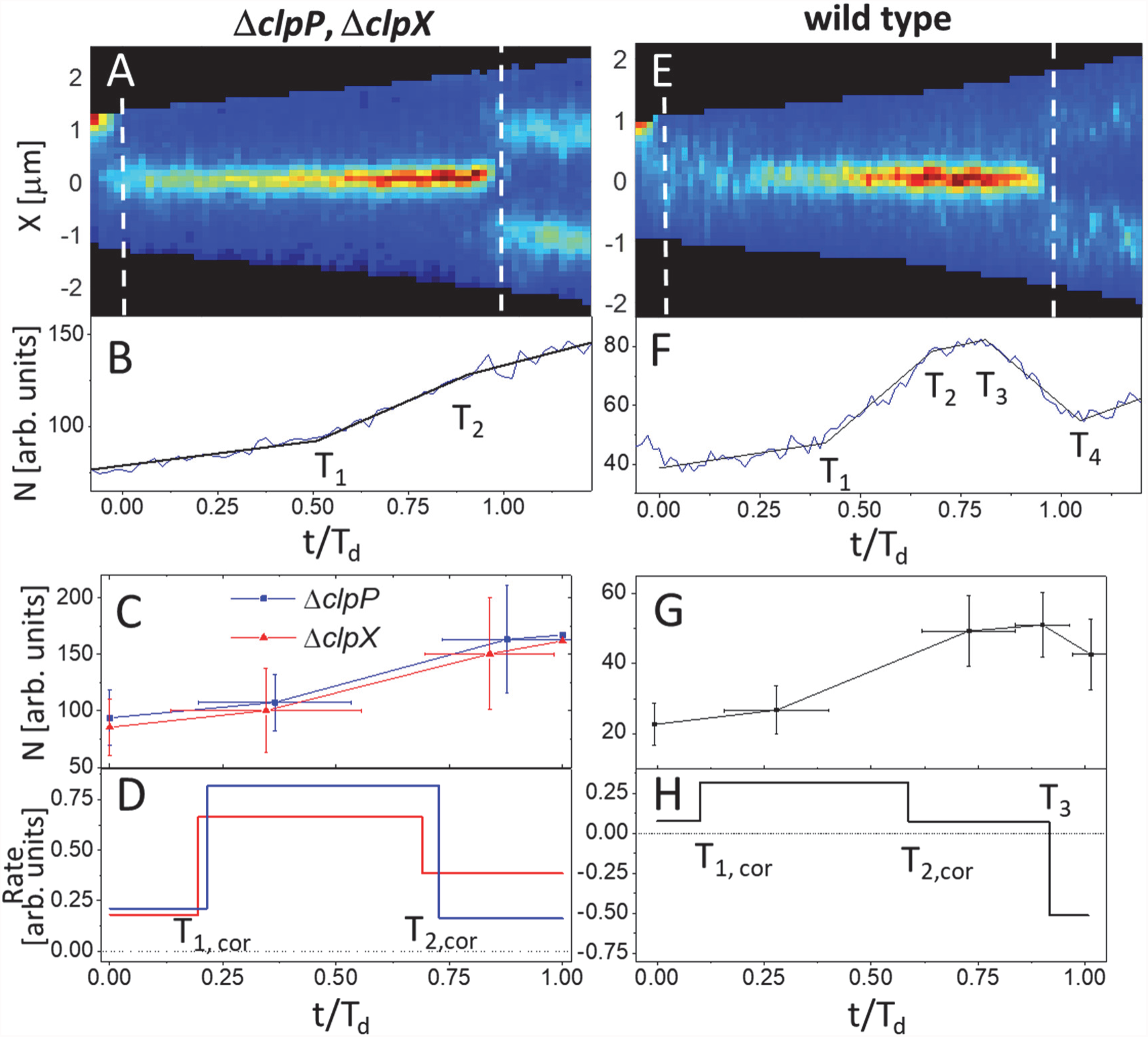
Rates of change in FtsZ numbers during cell cycle in ClpX and ClpP null mutant (left column) and WT (right column) strains in slow growth conditions. (A, E) Kymographs of FtsZ-mNG distribution along the long axes in representative cells. Panel (A) shows a Δ*clpP* cell. Vertical dashed lines mark cell divisions. (B, F) Number of fluorescent FtsZ-mNG as a function of the cell cycle time for the cells shown above. Solid black lines correspond to piecewise linear fit. T_1_ - T_4_ mark times when the slope of the curve changes. T_1_- upregulation of FtsZ synthesis; T_2_ - slow-down of the FtsZ synthesis; T_3_ - start of rapid FtsZ degradation, and T_4_ - end of rapid FtsZ degradation. (C, G) The fitting curves after averaging them over cell populations. N=35 for Δ*clpP*, N=32 for Δ*clpX* and N=53 for WT cells. Error bars are s. d. (D, H) rates of change in FtsZ-mNG numbers. The rates correspond to the slopes of the curves from the panel above. Times T_1,cor_ and T_2,cor_ have been corrected to account for the maturation of mNG.

The times T_1_ and T_2_ cited above did not include corrections for the maturation of the fluorescent reporter. To make this correction we measured maturation time of mNG in our cells using an approach described earlier (50) (see Materials and Methods). Measurements of WT cells in slow and fast growth conditions, and *ΔclpX* cells in slow growth conditions yielded all maturation half-lifetime of 25±3 min corresponding to (0.15±0.02)T_d_ (SI Fig. S2). Correcting for the maturation by subtracting maturation half-lifetime from the corresponding times (see SI *Modelling*, *Piecewise Linear Fitting*) yields corrected estimates T_1,cor_ = (0.2±0.2)T_d_ for the start and T_2,cor_= (0.7±0.15)T_d_ for the end of the faster synthesis period in *ΔclpP* and *ΔclpX* cells. Fig. 3D summarizes these results in terms of rate of change in FtsZ numbers during the cell cycle.

As expected from measurements of *ΔclpP* and *ΔclpX* strains, FtsZ synthesis rates also varied in WT cells during the cell cycle. In addition to upregulation in the synthesis rates, N vs t curves of individual cells also had a distinct period in the end of the cell cycle corresponding to rapid degradation of FtsZ (Fig. 3E-H). We included this period in the piecewise linear fits denoting the beginning of the period by T_3_ and its end by T_4_ (Fig. 3F, G). Including correction for the maturation time of mNG, FtsZ synthesis was upregulated in WT cells from T_1,cor_=(0.1±0.12)T_d_ to T_2,cor_=(0.6±0.1)T_d_ (mean±s.d.). During this period, the rate of FtsZ synthesis was about four times higher than in the beginning of the cell cycle (Fig. 3H) similar to the change observed in Δ*clpP* and Δ*clpX* cells. After the T_2_ point, the synthesis of FtsZ slows-down. From T_2,cor_= (0.6±0.1)T_d_ to T_3_ = (0.90±0.04)T_d_ FtsZ synthesis rates were comparable to those in the beginning of the cell cycle, i.e. from 0 to 0.1T_d_. The last period in the cell cycle from T_3_ = (0.9±0.04)T_d_ to T_4_=(1.0±0.05)T_d_ showed rapid decrease in FtsZ numbers. This period is clearly visible also in population average curves of Fig. 1E. Since the effect of degradation of the protein is immediate on the fluorescence no corrections were applied to the latter two times. The end of the degradation period coincided in good approximation with the end of the cell cycle. Fig. 3H summarizes the WT results in terms of rate of change in FtsZ numbers during the cell cycle.

It is noticeable that while the beginning (T_1_) and end (T_2_) of FtsZ synthesis upregulation showed substantial cell-to-cell variability, the beginning (T_3_) and end (T_4_) of fast degradation of FtsZ showed less stochasticity (Fig. 3G). This indicates that different types of regulation are involved in controlling synthesis and degradation. While synthesis is likely controlled at the transcriptional level and is known to have a high level of stochasticity (51) the same appears not to be the case for degradation.

We also attempted the same analysis for WT cells in fast growth conditions. Since degradation of FtsZ was not visible in most individual N vs t curves, only periods of slow and fast FtsZ synthesis were determined. The analysis indicated about two fold increase in FtsZ synthesis rate from T_1,cor_=0 to T_2,cor_=0.5T_d_ compared to the remainder of the cell cycle (SI Fig. S3). This difference is statistically significant in t-test (p<10^-5^ for equal slopes). However, taken that the maturation time of fluorescent reporter in these growth conditions is about a third of the doubling time, our analysis is expected to significantly underestimate differences in the rates and as such is not directly comparable to the values from slow growth conditions.

### Quantification of degradation rates in slow growth conditions

Using data from piecewise linear fits to *N* vs *t* curves from individual cells we next quantify the distribution of relative amounts of FtsZ degraded in the end of the cell cycle. We find a broad distribution of degraded amounts with an average of 22±14% (Fig. 4A). This amount is calculated relative to the number of FtsZs at end of the cell cycle. There appears to be no obvious correlation between the FtsZ amount at the end of the division and the amount of FtsZ degraded (SI Fig. S4). On average about the same number of FtsZ molecules are degraded in cells where FtsZ amounts differ by factor of two. The constant amount of FtsZ degraded can imply that the rate limiting step in degradation is not binding of ClpXP to FtsZ but the time it takes to degrade the protein. Based on single molecule measurements (52), typical time to degrade a single FtsZ-mNG molecule is about 10 sec. It is likely that this time exceeds the time needed for ClpXP and FtsZ to bind to each other.

**Figure 4.**
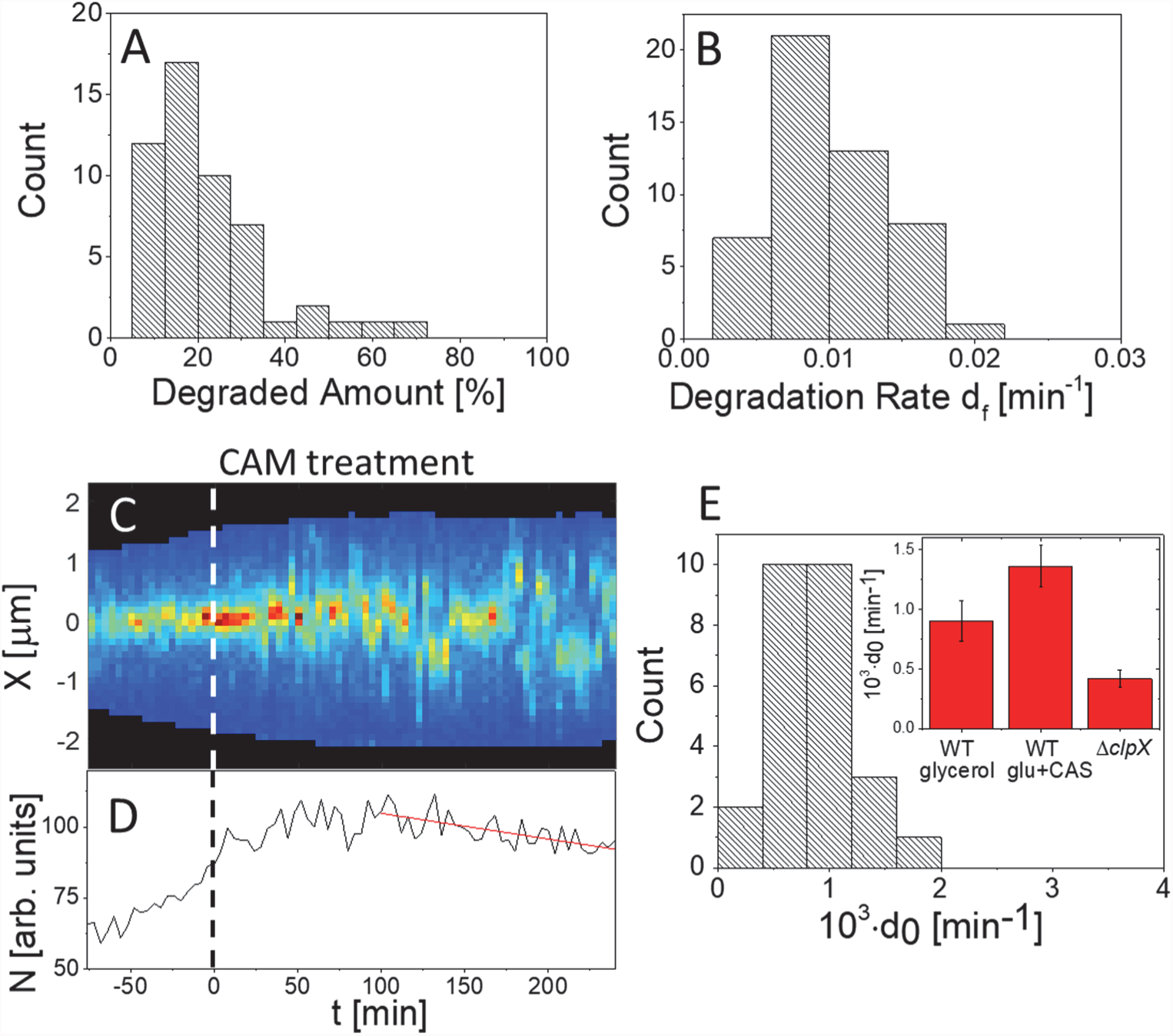
Degradation rates of FtsZ in the beginning and end of the cell cycle. (A) Degraded amount of FtsZ in the end of the cell cycle at the time when FtsZ disassociates from the Z-ring. The amount is relative to the amount at the end of cell division. (B) Degradation rate during this period. N=53. (C) Kymograph showing FtsZ-mNG distribution along the long axes of the cell before and during chloramphenicol treatment (CAM, 300 μg/ml). Drug is administered at time 0 min (vertical dashed line). (D) Amount of fluorescent FtsZ-mNG in this cell as a function of time. Basal degradation rate, *d_0_*, is determined from the linear fit to the curve (red solid line). (E) Distribution of *d_0_* in WT cell population in slow growth conditions (N=26). Inset shows the population average basal rate for WT cells in slow and fast growth conditions and for Δ*clpX* cells. The latter acts as a control. Error bars are s. d.

In addition to degraded amount, we also determined the time-average degradation rate at the end of the cell cycle (Fig. 4B). We found for the cell population the rate d_f_= 0.010±0.05 min^-1^. This rate is about factor two larger than the one found from bulk measurements in unsynchronized *E. coli* cultures (46). However, this difference should not be over interpreted because d_f_ depends on the protease concentration in the cell (see SI *Modelling*). The latter is likely to vary from strain to strain and from one growth condition to another.

We also determined the degradation rate of FtsZ during the remainder of the cell cycle. For that end we inhibited protein synthesis by chloramphenicol and observed decay in fluorescent signal of FtsZ-mNG (Fig. 4C,D). As described earlier, the fluorescent signal increases immediately after inhibition of protein synthesis because of maturation of the fluorescent label (50). To avoid this effect, we determined degradation rates after four mNG maturation half-lifetimes (100 min) from administering the drug (Fig. 4D). We found the average basal degradation rate d_0_=(0.9±0.4)·10^-3^ min^-1^ in slow and d_0_=(1.4±0.8)·10^-3^ min^-1^ in fast growth conditions (Fig. 4E). For Δ*clpX* strain the degradation rate constant was more than two times smaller d_0_=(0.4±0.2)·10^-3^ min^-1^. The apparent degradation rate of Δ*clpX* strain may reflect bleaching of mNG during the measurement or degradation of FtsZ by some other proteases. Subtracting values measured in Δ*clpX* strain from the WT one yields a ClpXP related degradation rate constant of d_0_=5·10^-4^ min^-1^ in WT cells. There is thus a 20-fold increase in ClpXP related degradation of FtsZ in the end of the cell cycle compared to basal rate.

### Z-ring dissociation triggers degradation of FtsZ by ClpXP

An order of magnitude increase in FtsZ degradation rate by ClpXP in the end of division invites the question, what triggers such a significant increase. To gain further insight we compared times when degradation of FtsZ (T_3_) starts to the times when FtsZ begins to dissociate from the Z-ring (T_dis_). We defined the latter as a time in the cell cycle when the total amount of FtsZ in the Z-ring starts to decrease (Fig. 5A). The comparison shows that the two times are highly correlated within the cell population (R=0.95) and essentially co-incident (Fig. 5B). The data also show that the total time it takes to dissociate FtsZ from the Z-ring does not depend whether ClpXP is present or absent (Fig. 5C). These data together indicate that FtsZ degradation does not drive Z-ring dissociation but is a consequence of it. One can furthermore conclude that the increase in the degradation rate is not triggered by an increase in ClpXP numbers but rather by its accessibility to FtsZ.

**Figure 5.**
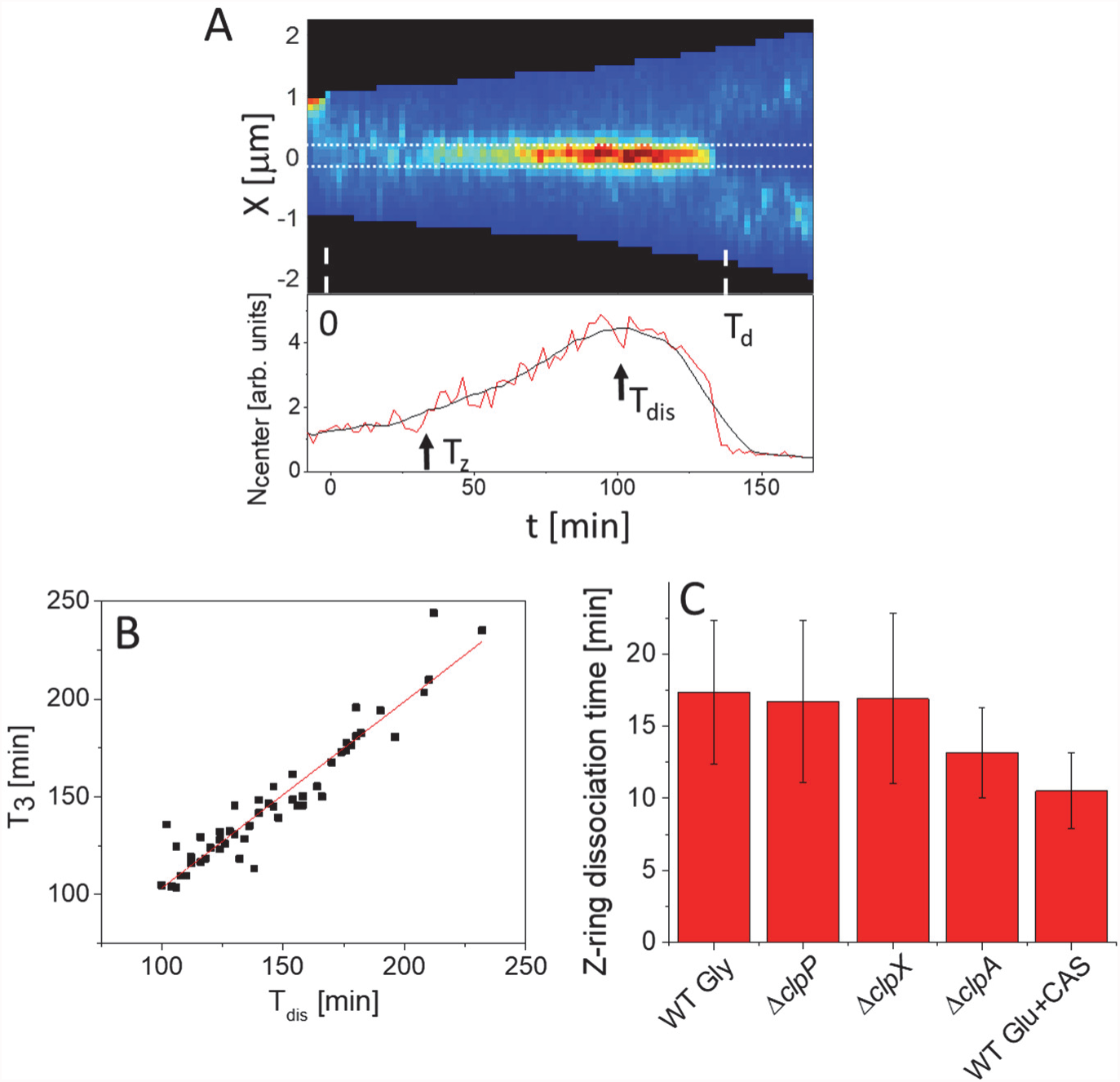
Rapid degradation of FtsZ is a consequence of the Z-ring dissociation. (A) Kymograph (top) and amount of fluorescent FtsZ-mNG at the cell center (bottom) in a representative cell. The cell center is defined as 300 nm wide band (indicated by horizontal dotted lines on kymograph). Fluorescent intensity after averaging by a running window is shown by a black line. T_z_ indicates start time and T_dis_ time when the Z-ring starts to dissociate. (B) Start time for FtsZ dissociation from constriction region (T_dis_) vs start time of rapid degradation of FtsZ (T_3_). (C) Total time for FtsZ to dissociate from the Z-ring (T_d_-T_dis_) in different strains and growth conditions. Error bars are s. d.

After FtsZ dissociates from the Z-ring it can be expected to be mostly monomeric. To determine if an increase in the numbers of monomeric FtsZ can explain its rapid degradation at the end of the cell cycle, we artificially induced dissociation of the Z-ring to monomeric FtsZ. For this purpose, we transformed the FtsZ-mNG strain with pA3 plasmid carrying inducible SulA (53). SulA binding to FtsZ in one to one stoichiometry lowers effective concentration of FtsZ in the cell so that it is below critical concentration for protofilament formation (53). SulA binding to FtsZ should leave the latter still accessible for ClpXP degradation because SulA does not bind to C-terminal domain of FtsZ, which contains the recognition elements for ClpX binding (46, 48). SulA upregulation and subsequent dissociation of the Z-ring, however, did not lead to any detectable degradation of FtsZ (SI Fig. S5A, B). Moreover, SulA upregulation did not change synthesis rate of FtsZ. In cells where SulA was upregulated in early stages of the cell cycle, FtsZ synthesis followed its well-defined pattern (cf. Fig. 3F) despite SulA expression (SI Fig. S5C, D). Consequently, the increase in monomer FtsZ concentration in the cytoplasm during Z-ring dissociation is not the reason for its rapid degradation at the end of the cell cycle.

### Correlations between FtsZ upregulation and Z-ring formation

Previously presented data (Fig. 3E) show that FtsZ synthesis rate increases 4-fold shortly after cell birth at 0.1T_d_ in slow growth conditions. We next investigate to what degree this upregulation correlates with the Z-ring formation. To answer this question we determined from kymographs, such as shown in Fig. 5A, the time when the Z-ring forms (T_z_). Note that in fast growth conditions, in almost all cells (94%), the Z-ring formed immediately after cell birth (SI Fig S6). However, in slow growth conditions there was a distinct delay between cell birth and the formation of the Z-ring (Fig. 6A). On average the Z-rings formed at T_z_=0.22T_d_ but the times varied considerably between individual cells (s.d. = 0.13T_d_). There was a similarly wide distribution in start times of FtsZ synthesis upregulation (s.d.=0.13T_d_, Fig. 6B). These times preceded on average about 0.1T_d_ (16 min) the times of Z-ring formation. At the level of individual cells the two times showed some level of correlation (R=0.67; Fig. 6C). In essentially all cells the Z-ring formed after FtsZ synthesis rate had increased. This indicates that upregulation of FtsZ levels is needed for formation of the Z-ring in slow growth conditions. Consistent with this conclusion, Z-rings formed earlier in cells where the amount of FtsZ was higher at cell birth (R=-0.43; Fig. 6D). At the same time the correlation between concentration of FtsZ at cell birth and timing of Z-ring formation were lower (R=-0.15; Fig. 6E) indicating that FtsZ numbers rather than its concentration is the factor controlling the Z-ring formation.

**Figure 6.**
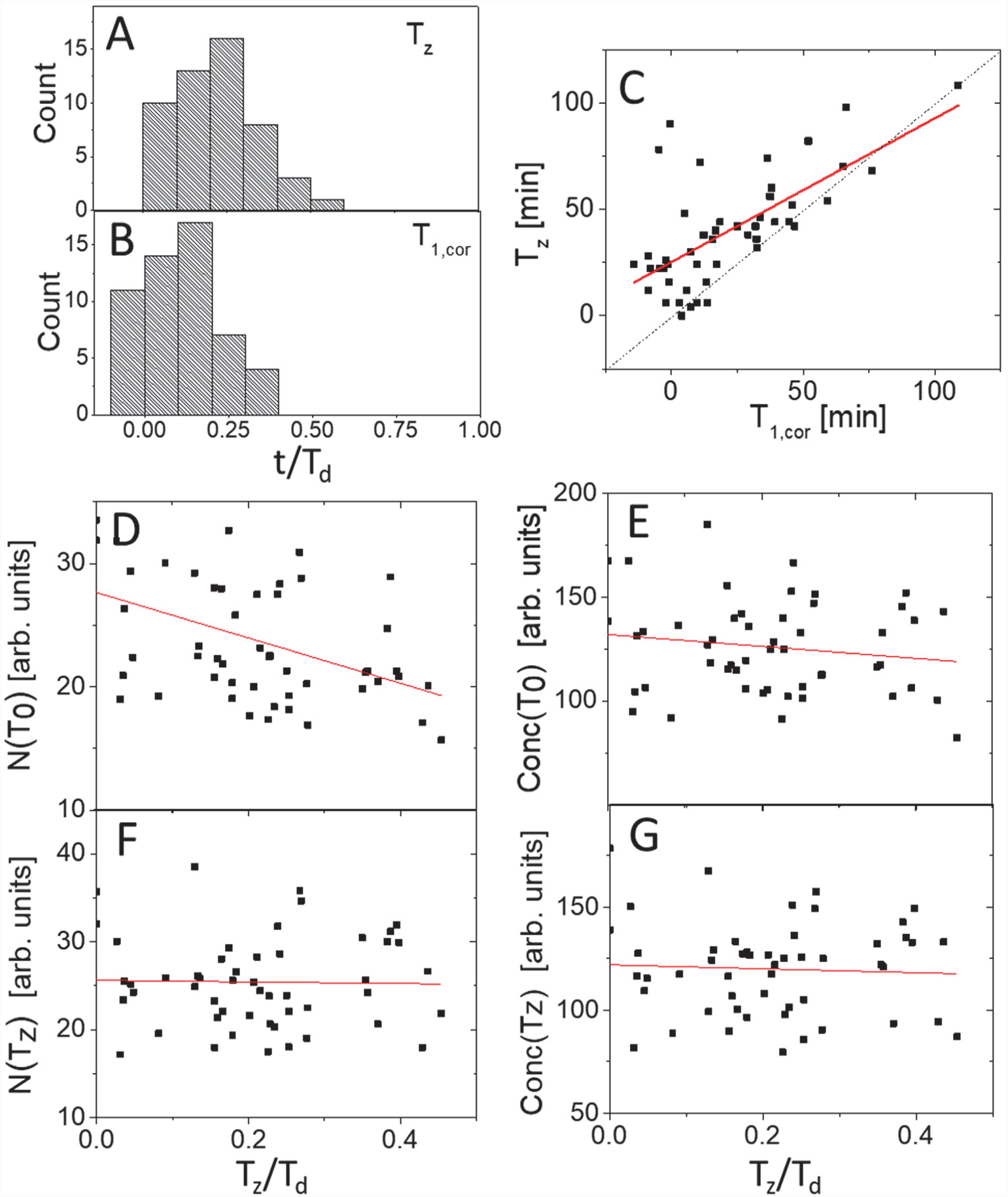
Link between upregulation of FtsZ synthesis and formation of the Z-ring in slow growth conditions. (A) Distribution of times when Z-ring forms (T_z_). (B) Distribution of times when FtsZ synthesis rate increases during the cell cycle (T_1,cor_). The times are corrected for mNG maturation time. (C) T_z_ vs T_1,cor_ for the same dataset. Dashed black line corresponds to T_z_ = T_1,cor_ and solid red line to linear fit (T_Z_ = −0.67T_1,cor_ + 25.2; R=-0.67). (D) Number of fluorescent FtsZ-mNG and (E) its concentration at cell birth (T_0_) vs. time when Z-ring forms. (F) Number of fluorescent FtsZ-mNG and (G) its concentration at the time of Z-ring formation vs. time when Z-ring forms. Solid lines are fits to the data. The fitting parameters are listed in SI Table S2. N=53.

At the time when the Z-ring formed, the amount of FtsZ in the cells (Fig. 6F) and its concentration (Fig. 6G) were about the same, and independent of the time the Z-ring formed (R=-0.02; R=-0.05, respectively). These data together show that cells with a low amount of FtsZ at birth synthesize more FtsZ to reach the level needed for Z-ring formation. However, the variation of both the amount of FtsZ and its concentration at the time of the Z-ring formation was large in cell population with a coefficient of variation CV = 20%. Large variation is indicative that FtsZ level is not the sole factor that controls the timing of the Z-ring formation. Altogether we conclude that FtsZ upregulation is necessary for the formation of the Z-ring in slow growth conditions but is not an immediate trigger for it.

## Discussion

We investigated changes in FtsZ numbers and concentration in slow and fast growth conditions in *E. coli*, and the effects these changes have on Z-ring formation. We found that in fast growth conditions the Z-ring formed in essentially all daughter compartments (94% of cells) concurrent with the dissociation of FtsZ from the constriction of the mother cell. This observation is consistent with previous reports (8, 54). Immediate formation of the Z-ring in fast growing cells indicates that there is no specific cell cycle control over this process. In contrast there is a distinct delay between cell birth and formation of the Z-ring in slow growth conditions. Even though the delay has large cell-to-cell variation (± 0.13T_d_), its presence indicates that there is a cell cycle based control over the Z-ring formation.

Our data shows that regulatory mechanisms controlling Z-ring formation in slow growth conditions includes a faster FtsZ synthesis rate in the first half and its rapid degradation in the last 10% of the cell cycle. The progression of these cell cycle events is quantitatively summarized in Fig. 7A, B. To further validate this model we compared it to numerically calculated FtsZ numbers and concentrations. The numerical calculation is based on experimentally determined rates including maturation rate of mNG (see SI *Modelling* for details). The only unknown parameter in these calculations is the conversion factor between one recorded fluorescent unit and emission from single FtsZ-mNG. Although we have estimated this conversion factor from photobleaching measurements, the uncertainties involved are large. Therefore we compare the experimental data and calculated curves after normalizing both by their maximum values. The normalized model and population average data show a good agreement (Fig. 7C, D). The two deviate from each other most significantly at time T_3_ when the degradation rate of FtsZ increases. The sharp cusp in the model is not present in population average data partially because of the spread of degradation start times in the population. It is also plausible that part of the discrepancy is caused by gradual increase in the degradation rate around time T_3_ instead of discontinuous change that we assumed in our analysis and model. Similarly to degradation rate one would also expect that the synthesis rate in individual cells shows a more complicated behavior than discontinuous change between constant levels at times T_1_ and T_2_. It is well-known that individual cell level protein synthesis show bursts of activity (55). These details go beyond our simplified model (Fig. 7A,B), which captures a population-average behavior on a coarse temporal resolution.

**Figure 7.**
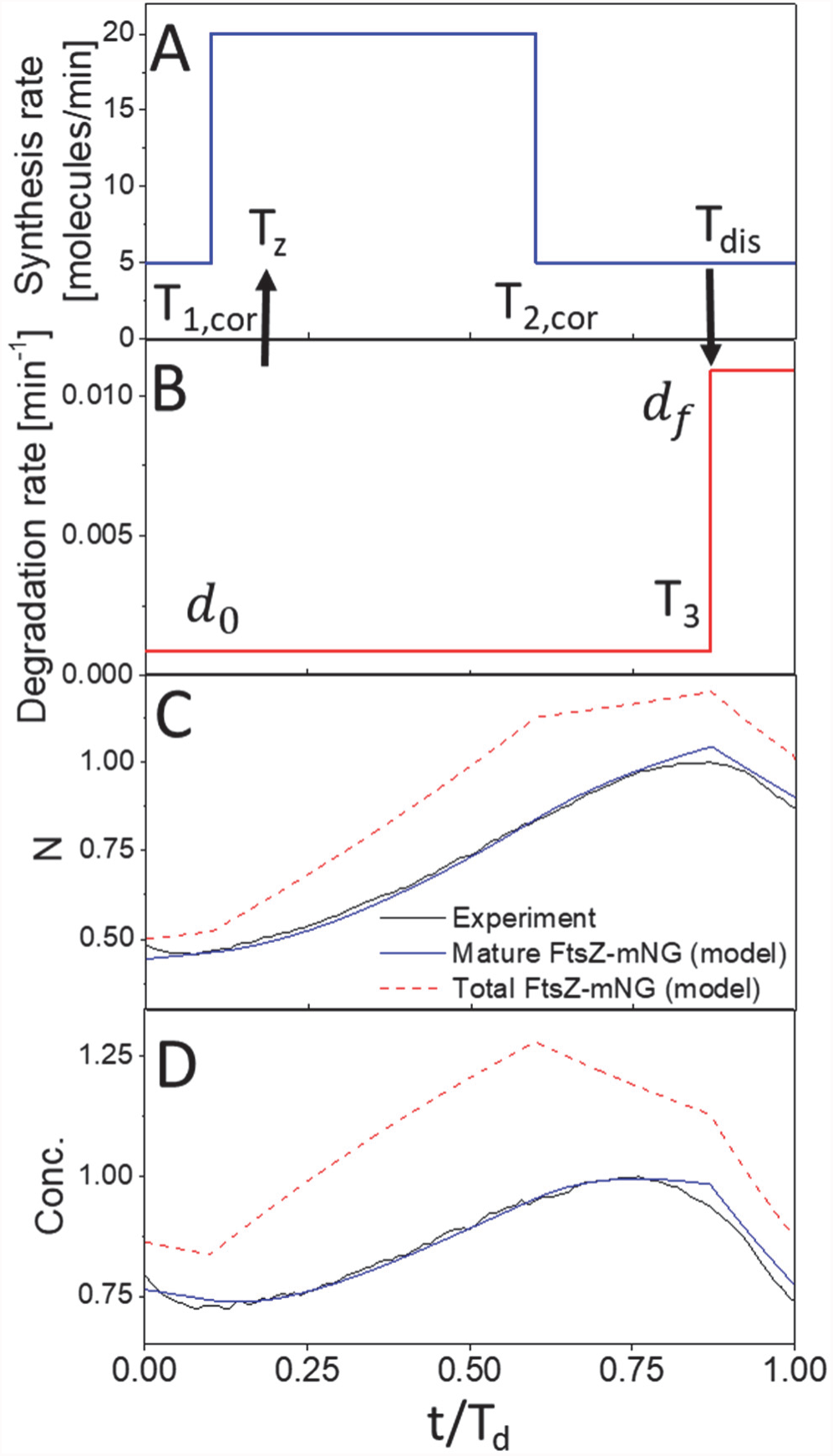
Comparing model and experiment. (A) Synthesis rate as function of time used in the model. The rate is 4 times the basal rate between T_1,cor_=0.1T_d_ and T_2,cor_=0.6T_d_ in accordance with the measurements. (B) Degradation rate as a function of time. Degradation rate increases from 0.001 min^-1^ to 0.010 min^-1^ at 0.9T_d_. (C) Comparison of number of FtsZ molecules in the cell. The numbers are normalized by the maximum number of mature FtsZ-mNG during the cell cycle. Solid black line corresponds to the experiment, blue line to mature, and dashed red line total FtsZ-mNG from the model. The additional model parameters not specified on panels (A) and (B) are T_d_=160 min and maturation rate kmat = 0.027 min^-1^. (D) Comparison of FtsZ concentrations. The concentrations have been normalized by maximum concentration of FtsZ-mNG during the cell cycle.

The FtsZ regulation cycle in slow-growing *E. coli* shows close resemblance to the one in *Caulobacter crescentus* swarmer cells. In *Caulobacter* swarmers, FtsZ synthesis is upregulated after transformation of a swarmer to a stalked cell occurs, and it is downregulated in the beginning and end of the cell cycle (56). Also, as we observed in slow growing *E. coli*, FtsZ is degraded in the end of the cell cycle (56, 57). In both organisms the degradation occurs via ClpXP based proteolysis, but in the case of *Caulobacter* ClpAP appears to play also a significant role (57). In our measurements the effect of ClpAP was modest (Fig. 3).

It has been found that upregulation of FtsZ synthesis in swarmer cells coincides with DNA replication period (56). Interestingly, we observe 4-fold upregulation of FtsZ synthesis in slow growing *E. coli* from 0.1T_d_ to 0.6T_d_. This period matches well with the period (0.14-0.51T_d_) reported for DNA replication in similar growth conditions (58). Increase in FtsZ mRNA levels upon initiation of DNA replication have been reported before in *E. coli* (33, 35). However, other studies have found maximal transcription levels around the middle of the cell cycle (32) or at the time of the division (34). While these different measurements lack yet consensus to support causative link between FtsZ upregulation and initiation of DNA replication in *E. coli*, a molecular level link between the two processes has been found in *C. crescentus* (56). This link is realized by the global cell cycle regulator CtrA that represses both the initiation of DNA replication and transcription of FtsZ (59).

Although FtsZ regulation during the cell-cycle is similar in *E. coli* and *C. crescentus* swarmer cells, the underlying molecular networks appear different. First, no global cell cycle regulator similar to CtrA is known to exist in *E. coli*. Second, in *Caulobacter* there is only a single promoter controlling FtsZ transcription (56) while in *E. coli* 7 different promoters are responsible for transcription of FtsZ (60). Of the seven the two proximal ones have been reported to lead to cell cycle dependent transcription of FtsZ (33), but in addition to promotors other regulatory elements such as RNA E cleavage sites, antisense RNA binding to *ftsZ* mRNA, and competition between ribosome binding sites on long polycistronic RNA could also play a role (60). Further work to understand cell cycle control on transcription FtsZ in *E. coli* is clearly warranted after nearly a two decade long pause in studies.

In addition to mechanisms controlling FtsZ synthesis, our work also raises question about how ClpXP is able to rapidly degrade FtsZ during the end of the cell cycle. The data show that dissociation of the Z-ring triggers degradation of FtsZ. After dissociation from the Z-ring FtsZ may lose specific protective interactions on its conserved C-terminal domain which has one of the two recognition sites for ClpX (46, 48). It has been reported before that one of FtsZ membrane linkers, ZipA, is able to protect FtsZ from degradation by ClpXP (61). Furthermore, it is possible that steric hindrance from the inner membrane reduces accessibility of ClpXP to the C-terminal tail of FtsZ and therefore protects it from degradation. In addition to FtsZ residing in the Z-ring, our data also show that monomeric FtsZ in the cytosol is not the preferred substrate for ClpXP. In light of these arguments we hypothesize that possible ClpXP substrates are FtsZ protofilaments or protofilament aggregates either free in the cytosol or partially attached to the membrane. This hypothesis is consistent with previous *in vitro* studies showing that the degradation rate of a filament or filament bundle is larger than monomeric FtsZ (46, 62).

To conclude, our data show that concentration of FtsZ oscillates in *E. coli* in a cell cycle specific manner in slow growth conditions. The lower FtsZ abundance at cell birth guarantees that the Z-ring does not form immediately at the beginning of the cell cycle. However, large variations in FtsZ numbers and concentrations at the time of Z-ring formation within the cell population indicate that some other factors beyond FtsZ can also play a role in triggering the Z-ring formation. Identifying these putative factors and establishing their link to the chromosome replication cycle remains an important avenue for future research.

## Materials and Methods

### Strains and growth conditions

All strains are derivatives of *E. coli* K-12 (Keio collection strain BW27783). The strains and plasmids are described in details in SI Table S1. All cells were grown and imaged in M9 minimal medium (Teknova) supplemented with 2 mM magnesium sulfate. In slow growth conditions 0.3% glycerol and in fast growth conditions 0.5% glucose supplemented with 0.2% casamino acids (ACROS Organics) were added to the media. During cell culturing on agar plates and in liquid media kanamycin (25 µg/ml) and chloramphenicol (25 µg/ml) were used as needed for all FtsZ-mNG expressing strains, and higher concentration of chloramphenicol (80 µg/ml) for the strain expressing FtsZ-GFP from the plasmid. No antibiotics were used during cell growth in microfluidic devices, except for a plasmid expressing FtsZ- GFP strain (chloramphenicol, 40 µg/ml). Cells were grown and imaged at 28 °C.

### Microfluidic chip fabrication

Soft-lithography of PDMS (polydimethylsiloxane) was used to fabricate microfluidic chips following previously described procedure (43). Briefly, silicon molds were fabricated combining e-beam and photolithography, and reactive ion etching. The mold defined dead-end channels, which were 0.8 μm wide, 1.1 μm high, and 20 μm long. PDMS (Sylgard 184, Dow Corning) was poured on passivated mold and baked for 15 min at 90 °C in a convection oven. Subsequently, the solidified PDMS layer was peeled from the mold and cut into pieces. Access holes to each piece, corresponding to one microfluidic device, were then punched using a biopsy needle. A PDMS piece was then plasma treated together with a clean #1.5 coverslip (Fisher Scientific). The two were bonded together after the treatment.

### Preparation and culturing *E. coli* in microfluidic devices

After overnight growth in liquid media,1 µg /ml of bovine serum albumin (BSA) was added to 1.5 ml of overnight cell culture (OD_600_> 0.4) and concentrated 100x by centrifugation. A 2-3 μl of resuspended culture was then pipetted into main flow channel of a microfluidic mother machine device. The cells were then allowed to populate the dead-end channels for about 1 hr. Once these channels were sufficiently populated, tubing was connected to the device, and the flow of fresh M9 medium with BSA (1 µg/ml) was started. The flow was maintained at 5 µl/min during the entire experiment by an NE-1000 Syringe Pump (New Era Pump Systems, NY). To ensure steady-state growth, the cells were left to grow in channels at least 14 hr before imaging started.

### Estimation of mNG maturation time by translational arrest with chloramphenicol

For chloramphenicol treatment extra tubing was connected to microfluidic chips that contained M9 media with 300 µg/ml chloramphenicol. Care was taken for the drug from the tubing not to reach the cells before treatment by backfilling part of the tubing with the regular medium. Once normal cell growth was verified in regular medium (2.5 hr), a separate syringe pump was started to circulate the medium with chloramphenicol (10 µl/min). Cells were imaged in the presence of chloramphenicol for 4 hr. After protein synthesis stopped as a result of chloramphenicol treatment, mNG fluorescence continued to increase due to maturation of the fluorophore. An increase of total fluorescence from mNG as a function of time allows determining the rate constant for maturation *kmat* from an exponential fit (SI Fig S2) (50).

### Inhibition of FtsZ polymerization by SulA expression

*E. coli* strain (MB43) expressing pA3 plasmid with the *sulA* gene under the control of *lac*-promoter (53) was used in these experiments. IPTG (80 μM), which was used for induction of the SulA, was delivered to cells following the same procedure as described for chloramphenicol.

### Microscopy

A Nikon Ti-E inverted fluorescence microscope with a 100X NA 1.40 oil immersion phase contrast objective and Perfect Focus system was used for imaging the bacteria in microfluidic channels. Fluorescence was excited by a 200W Hg lamp through ND4 and ND8 neutral density filters. Chroma 41001 filter-cube was used to record mNG and GFP images. Images were captured by an Andor iXon DU897 camera and recorded using NIS-Elements software. Power from Hg lamp was measured before time-lapse imaging using Thorlabs PM121D power meter. All strains were recorded at nominally the same illumination conditions except for the FtsZ-mNG strain in M9 glycerol, which exposure was increased twice due to its lower level of FtsZ.

### Data analysis

Matlab, along with the Image Analysis Toolbox and DipImage Toolbox, (http://www.diplib.org/) was used for image analysis. In all analysis of time-lapse recordings, corrections to subpixel shifts between different frames were applied first. These shifts were determined by correlating phase contrast images in adjacent frames. The cells were then segmented based on phase contrast images using a custom Matlab script. The script first identified the coordinates of the two poles of each cell. These coordinates defined the long axes of the cell and yielded cell length. Intensity line profiles along the axes were then determined by integrating parallel line profiles along this centerline and subtracting the background. These profiles were plotted as kymographs for each cell. Segmentation routine allowed for joining daughter cells after their division. The total intensity from the profiles integrated over cell length is proportional to the number of fluorescent FtsZ-mNG in the cell (N). Concentration was calculated by dividing total intensity by the cell length assuming the same widths for all the cells.

Piecewise linear fits to total fluorescence curves were carried out using a custom Matlab script. Number of different segments for Δ*clpP* and Δ*clpX* strains was four and for WT strains six. The end point intensities and times for each segment were treated as fitting parameters. The exponential fits to find maturation rate of mNG and linear fits to find basal rate of FtsZ degradation were carried out in Origin 2016 software. Further discussion on accuracy of this procedure is given in SI *Modelling*.

## Acknowledgements

The authors thank Harold Erickson, Jie Xiao and Alex Dajkovic for strains and plasmids, Da Yang and Scott Retterer for help in microfluidic chip making, and Harold Erickson and Maxim Lavrentovich for valuable comments. Authors acknowledge technical assistance and material support from the Center for Environmental Biotechnology at the University of Tennessee. A part of this research was conducted at the Center for Nanophase Materials Sciences, which is sponsored at Oak Ridge National Laboratory by the Scientific User Facilities Division, Office of Basic Energy Sciences, U.S. Department of Energy. This work has been supported in part by NSF research grant MCB-1252890.

